# Transcriptional states of retroelement-inserted regions and KRAB zinc finger protein association regulate DNA methylation of retroelements in human male germ cells

**DOI:** 10.1101/2021.05.19.444783

**Authors:** Kei Fukuda, Yoshinori Makino, Satoru Kaneko, Yuki Okada, Kenji Ichiyanagi, Yoichi Shinkai

## Abstract

DNA methylation, repressive histone modifications, and PIWI-interacting RNAs are essential for controlling retroelement silencing in mammalian germ lines. Dysregulation of retroelement silencing is associated with male sterility. Although retroelement silencing mechanisms have been extensively studied in mouse germ cells, little progress has been made in humans. Here, we show that the Krüppel-associated box domain zinc finger proteins (KRAB-ZFPs) are associated with DNA methylation of retroelements in human primordial germ cells (hPGCs), and hominoid-specific retroelement SINE-VNTR-*Alus* (SVA) is subjected to transcription-directed *de novo* DNA methylation during human spermatogenesis. Furthermore, we show that the degree of *de novo* DNA methylation in SVAs varies among human individuals, which confers a significant inter-individual epigenetic variation in sperm. Collectively, our results provide potential molecular mechanisms for the regulation of retroelements in human male germ cells.

## INTRODUCTION

Transposable elements comprise more than 40% of most extant mammalian genomes (Lander et al. 2001). Among these, certain types of transposable elements called retroelements, including short/long interspersed elements (SINEs/LINEs), endogenous retroviruses (ERVs), and hominoid-specific retrotransposons SINE-VNTR-*Alus* (SVA) are still active and capable of transposition (Ostertag et al. 2003; Huang et al. 2012; Maksakova et al. 2013). As retrotransposons cause genome instability, insertional mutagenesis, or transcriptional perturbation and are often deleterious to host species (O’Donnell and Burns 2010), multiple defense mechanisms have been established against transposition. The first line of defense is transcriptional silencing of integrated retroelements by chromatin modifications, such as DNA methylation and histone H3 lysine 9 (H3K9) methylation (Goodier 2016; Fukuda and Shinkai 2020). Most retroelement families are bound by Krüppel-associated box domain zinc finger proteins (KRAB-ZFPs), which coevolve to recognize specific retroelement families (Jacobs et al. 2014; Wolf et al. 2015; Imbeault et al. 2017). KRAB-ZFPs repress retroelements by recruiting KAP1/TRIM28 (Sripathy et al. 2006) and other repressive epigenetic modifiers (Schultz et al. 2001; Schultz et al. 2002).

Restricting retroelements is especially important for germ cells because of their transmission to the next generation. During embryonic development, primordial germ cells (PGCs) undergo epigenetic reprogramming, characterized by both DNA demethylation and global resetting of histone marks in mice and humans (Seki et al. 2007; Seisenberger et al. 2012; Kobayashi et al. 2013; Gkountela et al. 2015; Guo et al. 2015; Tang et al. 2015). A subset of young retroelements escape from this global DNA demethylation event in PGCs, which might be required for retroelement silencing (Seki et al. 2007; Seisenberger et al. 2012; Kobayashi et al. 2013; Gkountela et al. 2015; Guo et al. 2015; Tang et al. 2015). H3K9me3 mediated by SETDB1 is enriched in DNA demethylation-resistant retroelements in mouse PGCs (Liu et al. 2014). As SETDB1 regulates DNA methylation of a subset of retroelements (Matsui et al. 2010; Rowe et al. 2013), and it is recruited to the retroelements via interaction with KRAB-ZFPs; it is speculated that SETDB1/KRAB-ZFPs contribute to DNA demethylation resistance in PGCs. In contrast to the extensive DNA hypomethylation in PGCs, genomic DNA of sperm is highly methylated in both humans and mice (Molaro et al. 2011; Kobayashi et al. 2013; Hammoud et al. 2014; Okae et al. 2014). Retroelements are also subjected to *de novo* DNA methylation during spermatogenesis in mice via the PIWI/piRNA-pathway and others (Aravin et al. 2008; Inoue et al. 2017). Epigenetic alterations in retroelements and dysfunction of retroelement silencing pathways in male germ cells are associated with male sterility linked to azoospermia (Bourc’his and Bestor 2004; Aravin et al. 2007; Carmell et al. 2007; Heyn et al. 2012; Urdinguio et al. 2015). In addition, epigenetic alterations of retroelements in male germ cells can potentially be transmitted to the next generations and change their phenotypes (Rakyan et al. 2003; Daxinger et al. 2016). Therefore, deciphering the regulatory mechanisms of retroelements in germ cells contributes to understanding sterility and transgenerational epigenetic inheritance. Despite extensive studies that have been conducted to understand DNA methylation mechanisms in mouse spermatogenesis, only limited progress has been achieved in humans.

In this study, we performed an integrative analysis of three sets of previously reported data: whole-genome bisulfite sequencing (WGBS) data of human PGCs (hPGCs) and sperm, transcriptome of human male germ cells, and comprehensive human KRAB-ZFPs ChIP-exo data. From this analysis, we revealed that KRAB-ZFPs are associated with DNA demethylation resistance of retroelements in hPGCs. We also found that *de novo* DNA methylation patterns in spermatogenesis vary among L1, LTR, and SVA retroelements. Notably, we found that SVAs are subjected to *de novo* DNA methylation by the transcription-directed DNA methylation machinery. Interestingly, the effectiveness of the machinery also varies among human individuals, with SVAs being one of the major sources of epigenetic variations in sperm.

## RESULT

### Transposable elements showing DNA demethylation resistance in hPGCs

To reveal the determinants of DNA demethylation resistance in hPGCs, we reanalyzed publicly available WGBS data for male hPGCs (Guo et al. 2015). As the global erasure of DNA methylation was mostly completed at 19 weeks of gestation (Fig. 1A), we further analyzed the DNA methylation status of full-length transposable elements in male hPGCs at 19 weeks of gestation to search for retroelements that display demethylation resistance. Typically, we focused on full-length copies of retroelements to analyze DNA methylation for at least 30 copies. Among the different retroelements analyzed, primate-specific retroelement families L1PA, SVA, and LTR12 showed high DNA methylation status (Fig. 1B). In the SVA family, SVA_A, which was the oldest SVA type and emerged 13–14 million years ago (Mya), showed the highest DNA methylation levels than other SVA types, including currently active SVA_E/F (Fig. 1C). In the L1 family, L1PA3-5, which were moderately young and emerged 12–20 Mya, showed higher methylation levels than the older L1 types (L1PA5–8) and younger L1 types, including currently active L1 (L1HS) (Fig. 1D). LTR12 (also known as HERV9 LTR) is not currently active, and all LTR12 types were highly methylated (Fig. 1E). Thus, it seemed that young but not active ones among L1PA, SVA, and LTR12 types tended to show DNA demethylation resistance in hPGCs. In addition, DNA methylation levels of each retroelement type were highly variable among full-length copies (Fig. 1C-E), which prompted us to identify the potential determinants of DNA sequences for DNA demethylation resistance by comparing DNA sequences of retroelement copies. To this end, we classified each retroelement copy according to their DNA methylation levels as follows: low < 20%, 20% ≤ medium < 60%, high ≥ 60%. From this classification, we observed that both the “High” and “Low” classes of copies exist in highly methylated retroelement types in hPGCs such as SVA_A, L1PA3, and LTR12C (Fig. 1F-H). Next, we examined the differences between the groups.

**Fig. 1.**
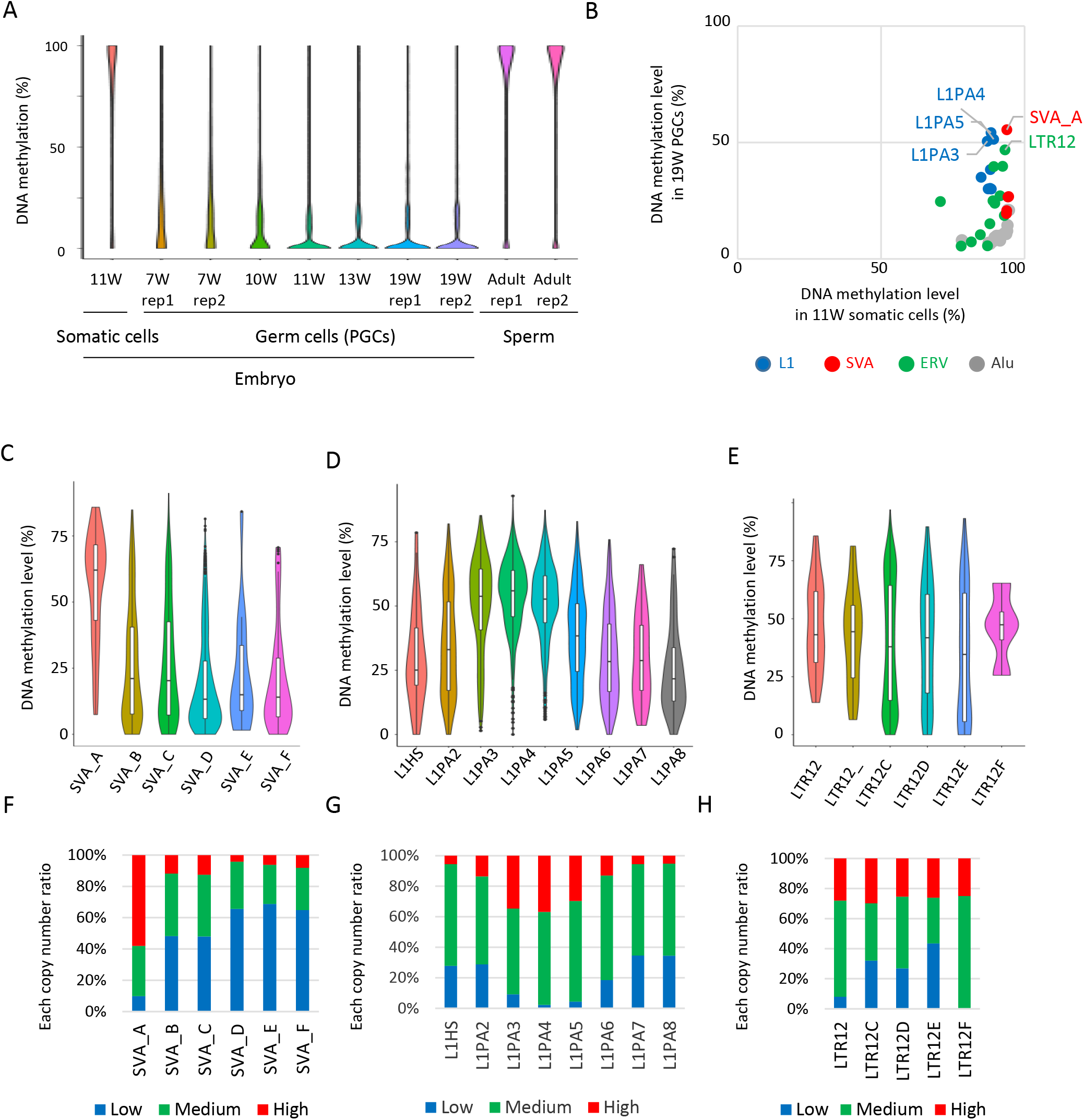
Retroelements showing DNA demethylation resistance (A) Violin plots showing DNA methylation levels of each CpG site during human male germ-cell development. DNA demethylation was almost completed at 19 weeks of gestation. (B) Scatter plots showing average DNA methylation level of each retroelement type between somatic cells and male hPGCs at 19 weeks of gestation. Only full-length copies were used for this analysis, and only retroelement types with >= 30 full-length copies were shown. Each plot was colored according to its retroelement family (Red: SVA, Blue: L1, Green: LTR, Gray: Other). (C-E) Violin plots showing DNA methylation level of each retroelement type in hPGCs at 19 weeks of gestation. (F-H) Bar graphs showing the fraction of “Low”, “Medium” and “High” methylated class of each retroelement type in male hPGCs at 19 weeks of gestation.

### The presence of ZNF28 and ZNF257 binding motifs are correlated with demethylation resistance in SVA_A

KRAB-ZFPs are pivotal factors for retroelement silencing by recruiting KAP1 and SETDB1. To investigate whether KRAB-ZFPs are involved in DNA demethylation resistance of SVAs, we analyzed the frequency of SVA_A copies with previously reported 236 KRAB-ZFP peaks (Imbeault et al. 2017). We found that ZNF257 and ZNF28 binding was correlated with the DNA methylation of SVA_A (Fig. 2A). Although ZNF611 and ZNF91 have been reported to interact with SVAs in mESCs (Jacobs et al. 2014; Haring et al. 2021) and those bindings were confirmed on the SVA_A copies (Fig. 2A), the interaction was not correlated with DNA methylation states of SVAs in male hPGCs at 19 weeks of gestation (Fig. 2A). More than 53.8% of “High” or “Medium” SVA_A elements were bound by either ZNF257 or ZNF28, while no “Low” SVA_A was bound (Fig. 2B). Enrichment of ZNF257 and ZNF28 in SVA_A was positively correlated with DNA methylation (Fig. 2C), and both ZNF257 and ZNF28 showed the highest enrichment of SVA_A in the SVA family (Fig. 2D). The SVA element contains a region of variable number tandem repeats (VNTRs) in the middle part. SVA_A contains one type of VNTR (VNTR1), while other SVA classes possess two types of VNTRs (VNTR1 and VNTR2) (Fig. 2F). ZNF257 and ZNF28 binding motifs, which were predicted by HOMER (Heinz et al. 2010) (Fig. 2E), were located in VNTR1 (Fig. 2F). The number of ZNF257 and ZNF28 binding motifs along SVAs was the largest in SVA_A (Fig. 2F), which was correlated with the largest copy number of VNTR1 in SVA_A among SVA classes (Fig. 2G). The VNTR1 copy number was also highly variable among SVA_A copies (Fig. 2G), and DNA methylation of SVA_A was positively correlated with the VNTR1 copy number (Fig. 2H). The ZNF257/28 motif number was also correlated with the DNA methylation status of SVA_A (Fig. 2I). These data indicate that a high number of ZNF257 and ZNF28 binding motifs, which is caused by high VNTR1 copy number, enhances the enrichment of these KRAB-ZFPs, and maintains DNA methylation of SVA_A during hPGC development. Although ZNF257 and ZNF28 protein expression during male hPGCs development are not examined yet, at least their transcript were detected in male hPGCs (RPKM of *ZNF257* and *ZNF28* were 3.49 and 0.66, respectively) (Gkountela et al. 2015).

**Fig.2.**
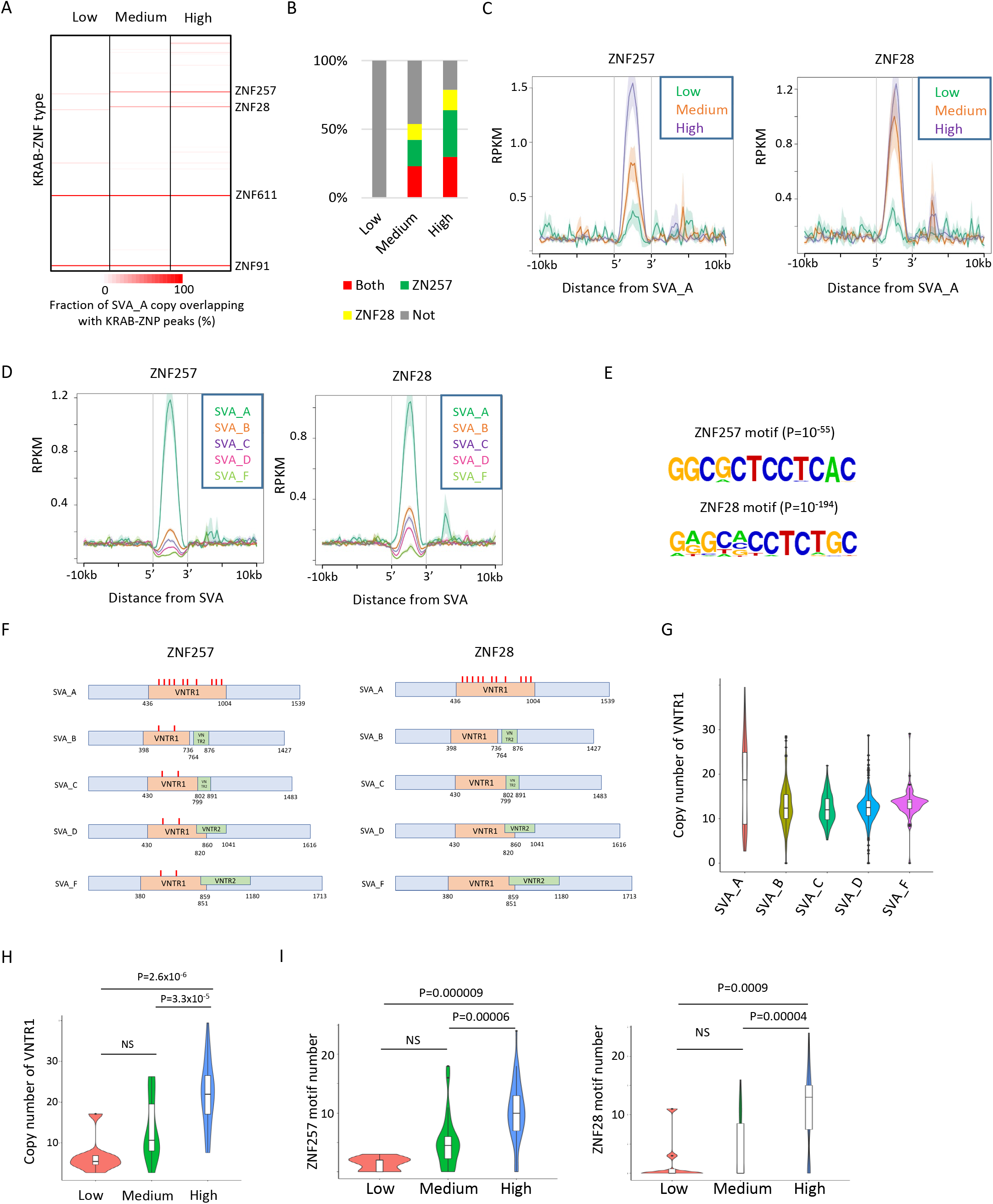
Identification of KRAB-ZFPs associated with DNA demethylation resistance in SVAs (A) Heatmap showing the fraction of SVA_A copies which overlaps of KRAB-ZFP peaks. ZNF257 and ZNF28 peaks were more frequently overlapped with “Medium” and “High” methylated SVA_A than “Low” methylated SVA_A. For this analysis, publicly available ChIP-exo data from 236 human KRAB-ZFPs in HEK293T cells (Imbeault et al. 2017) were used. (B) Bar graphs showing the fraction of SVA_A copies with ZNF257 and ZNF28 peaks. (C) Enrichment of ZNF257 and ZNF28 on SVA_A classified by DNA methylation levels in male hPGCs at 19 weeks of gestation. (D) Enrichment of ZNF257 and ZNF28 on each SVA type. (E) Sequence logo of ZNF257 and ZNF28 binding motifs. (F) Position of ZNF257 and ZNF28 binding motifs along SVA consensus sequences. VNTR1 and VNTR2 is composed of multiple copy number of tandem repeats, and the copy number of these VNTR is highly variable among SVA copies. Both ZNF257 and ZNF28 binding motifs were found within VNTR1 of SVAs. (G) Violin plots showing copy number of VNTR1 of each SVA type. (H) Violin plots showing VNTR1 copy number of SVA_A classified by its DNA methylation status in male hPGCs at 19 weeks of gestation. (I) Violin plots showing the number of ZNF257 and ZNF28 motifs in SVA_A classified by DNA methylation status in male hPGCs at 19 weeks of gestation. P-value was calculated by Tukey’s test.

### The presence of the ZNF649 binding motif is correlated with demethylation resistance in L1s

We also analyzed the association of KRAB-ZFP binding motifs and DNA methylation status of L1s and LTR12s in hPGCs. Consistent with previous reports that ZNF649 and ZNF93 bind L1s (Jacobs et al. 2014; Cosby et al. 2019), ZNF649 and ZNF93 peaks were frequently found in L1PA2–6 and L1PA3–6, respectively (Fig. 3A), and these two KRAB-ZFPs were enriched at the 5’ terminus of L1 sequences (Fig. 3B). The frequency of L1 copies with ZNF649 or ZNF93 peaks correlated with the DNA methylation levels of L1s (Fig. 3C). Therefore, these two KRAB-ZFPs are candidate factors for the DNA demethylation resistance of L1s. Similar to the SVA_A case described above, the presence of ZNF649 or ZNF93 binding motifs (Fig. 3D) was correlated with DNA methylation levels (Fig. 3E). The correlation between the binding motif and DNA methylation was stronger in ZNF649 than in ZNF93 (Fig. 3E); ZNF93 was not bound to L1PA2, which could be methylated in hPGCs (Fig. 3A, B), and it was reported that ZNF649 was expressed in male hPGCs, but ZNF93 not (Fig. 3F) (Gkountela et al. 2015). These data suggest that ZNF649 plays a more central role in the DNA demethylation resistance of L1s in hPGCs than ZNF93. The ZNF649 binding motif was located at the 5' UTR of L1s (Fig. 3G), consistent with the enrichment of ZNF649 in that region (Fig. 3B). Enrichment of ZNF649 in L1s was decreased in L1PA2 and almost lost in L1HS (Fig. 3B). Consistent with the reduced ZNF649 enrichment, base substitution at the fifth position of the ZNF649 binding site occurred in the consensus sequences of L1HS (Fig. 3G). As the fifth position of the ZNF649 binding site (T) tended to be conserved in highly methylated L1 copies (Fig. 3H), this position of T might be required for ZNF649 binding to L1. Unlike SVAs and L1s, no KRAB-ZFPs showed a correlation between their enrichment and DNA methylation of the LTR12 family in hPGCs (Fig. 3I). However, phylogenic analysis of LTR12C copies showed that highly-methylated LTR12C copies were genetically separated from low-methylated copies (Fig. 3J). Thus, the DNA demethylation resistance of LTR12C was also genetically determined.

**Fig. 3.**
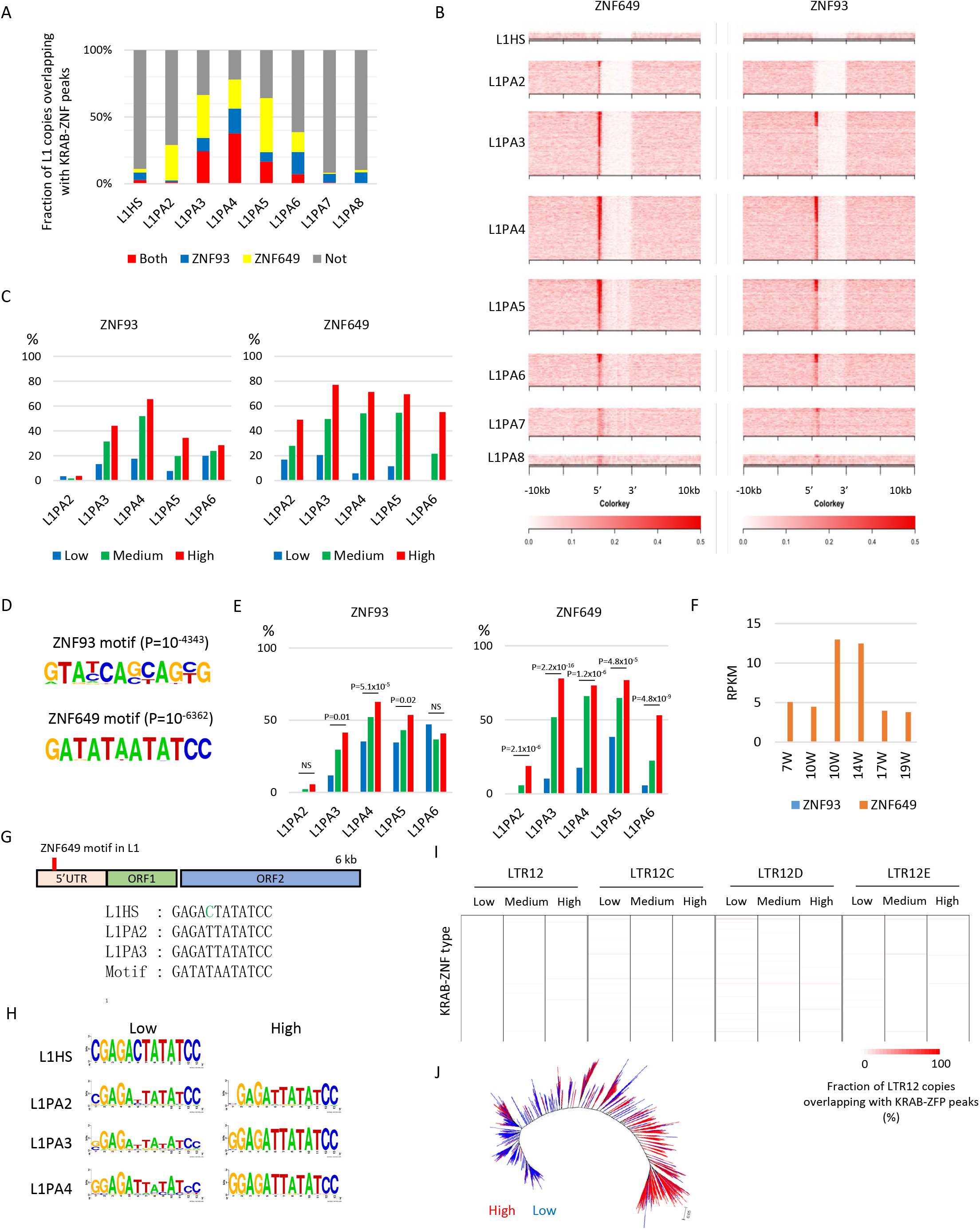
Identification of KRAB-ZFPs associated with DNA demethylation resistance in L1 (A) Bar graphs showing the fraction of full-length L1 copies with ZNF93 and ZNF649 peaks. (B) Heatmaps showing enrichment of ZNF649 and ZNF93 along full-length L1 copies. ZNF649 binds 5’ regions of L1PA2∼PA8, while ZNF93 binds the 5’ regions of L1PA3∼PA8. (C) Bar graphs showing the fraction of L1 copies with ZNF93 and ZNF649 peaks. L1 copies were classified by their type and DNA methylation levels in male hPGCs at 19 weeks of gestation. (D) Sequence logo of ZNF93 and ZNF649 binding motifs. (E) Bar graphs showing the fraction of L1 copies with ZNF93 and ZNF649 binding motifs. The presence of ZNF93 and ZNF649 binding motifs was correlated with higher DNA methylation of L1 in male hPGCs at 19 weeks of gestation. P-value was calculated by Hypothesis Testing for the Difference in the Population Proportions using a function of prop.test byR. (F) Expression of ZNF93 and ZNF649 during male hPGC development. (G) Comparison of sequences of ZNF649 binding sites among L1 types. L1HS lost the ZNF649 motif by a base substitution. (H) Comparison of sequences at ZNF649 binding sites between low and high methylated L1. (I) Heatmap showing the fraction of LTR12C copies, which overlap of KRAB-ZFP peaks. No KRAB-ZFP peak was correlated with DNA methylation levels of LTR12C in hPGCs. (J) Phylogenetic analysis of LTR12C copies classified by DNA methylation levels in hPGCs. Low and High methylated LTR12C copies were colored blue and red, respectively.

### Mode of DNA methylation acquisition during spermatogenesis is different among retroelement types

To investigate how the DNA methylation status of retroelements changes during spermatogenesis, we compared DNA methylation levels of each retroelement copy in male hPGCs at 19 weeks of gestation to those in adult sperm. For this analysis, publicly available human sperm WGBS data from two donors were used (Hammoud et al. 2014). The dynamics of DNA methylation in retroelements during spermatogenesis are vastly different among retroelement types and individuals. The majority of L1 copies acquired DNA methylation during spermatogenesis in both individuals, while LTR12C copies tended to maintain their DNA methylation status in hPGCs during spermatogenesis (Fig. 4A). A substantial difference between individuals was found in SVAs. The majority of SVA copies acquired DNA methylation during spermatogenesis in sperm donor 1 but not in sperm donor 2 (Fig. 4A). To show these trends more efficiently, we classified retroelement copies according to DNA methylation levels in sperm (common high: >60% in both donors, high and low: >60% in donor1 and <20% in donor2, common low: <20% in both donors). The majority of low-methylated L1 copies in hPGCs were highly methylated in sperm from both donors (Fig. 4B). In contrast, most LTR12C/D copies tended to maintain the PGC DNA methylation status during spermatogenesis (Fig. 4C). Among the SVA types, SVA_A tended to show high DNA methylation levels in both sperm donors, while other SVA types showed variable methylation levels between sperm donors (Fig. 4D), especially in the SVA copies with low DNA methylation levels in hPGCs (Fig. 4E).

**Fig. 4.**
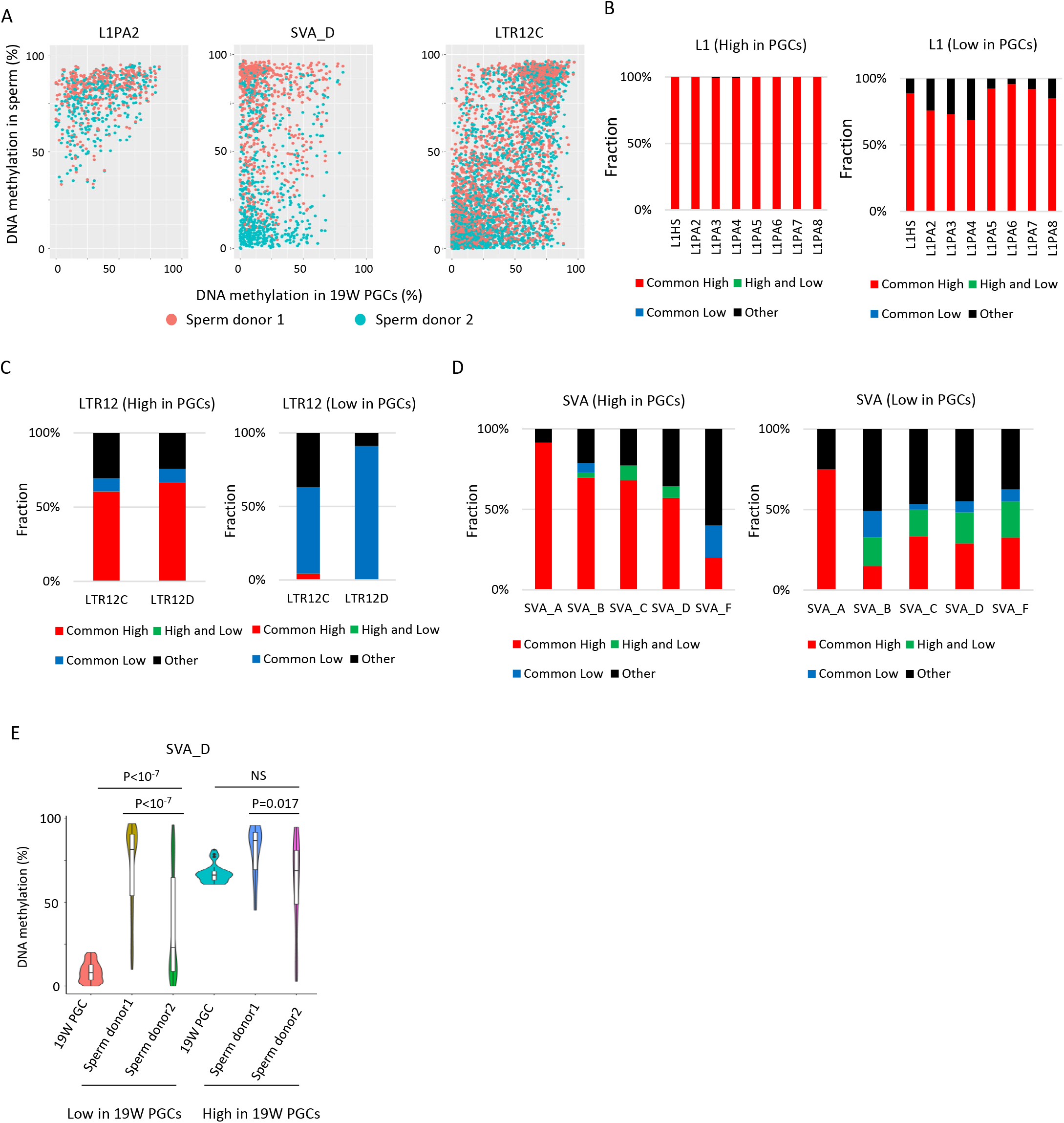
DNA methylation dynamics of retroelements during human spermatogenesis (A) Scatter plots showing DNA methylation levels of each retroelement copy in male hPGCs at 19 weeks of gestation and sperm. WGBS data from two sperm donors (Hammoud et al. 2014) were used for this analysis. Donor1 and donor2 were colored by orange and cyan, respectively. (B-D) Bar graphs showing the fraction of groups determined by DNA methylation patterns in two sperm donors in L1 (B), LTR12 (C), and SVA (D). Bar graphs were also separated by DNA methylation levels (high or low) in male hPGCs at 19 weeks of gestation. (E) Violin plots showing DNA methylation levels of SVA_D copies in male hPGCs at 19 weeks of gestation, sperm donor1, and sperm donor2. The violin plots were also separated by DNA methylation levels of SVA_D copies in male hPGCs at 19 weeks of gestation. Although hypomethylated SVA_D copies in male hPGCs at 19 weeks of gestation acquired DNA methylation during spermatogenesis, the degree of DNA methylation increase was significantly different between sperm donors. P-value was calculated by Tukey’s test.

### The degree of DNA methylation acquisition during spermatogenesis is different among SVA copies

Although the DNA methylation status of SVAs was highly variable between sperm donors, some SVA copies acquired DNA methylation or maintained a low-methylated state during spermatogenesis in both sperm donors (Fig. 4D). Thus, we can address how SVAs gain DNA methylation during spermatogenesis by comparing SVAs acquiring DNA methylation in both sperm donors (“Low” in hPGCs and “Common High” in sperm) to SVAs maintaining hypomethylation in both sperm donors (“Low” in PGCs and “Common Low” in sperm). Phylogenetic analysis of “Common Low” and “Common High” SVA_B copies showed that these two classes of SVA copies were not genetically separated (Fig. 5A), which suggests that acquisition of DNA methylation in SVAs during spermatogenesis is not genetically determined. The presence of transcription-directed retroelement silencing mechanisms, such as the PIWI/piRNA pathway (Watanabe et al. 2018), prompted us to investigate the correlation between the genomic distribution of SVA copies and DNA methylation. Approximately half of SVA_B–F were inserted in the gene body, most of which were in the antisense direction (Fig. 5B). Interestingly, “Common High” SVA_B–F copies were enriched in the gene body with antisense direction, and “Common Low” SVA_B–F copies were depleted from the gene body (Fig. 5B). We reanalyzed previously reported single-cell RNA-seq data in human testes (Sohni et al. 2019) to investigate the expression status of genes with SVAs in the antisense direction. Genes with “Common High” SVA_B–F copies tended to show a higher expression in spermatogonial stem cells than genes with “Common Low” (Fig. 5C). Thus, SVAs are methylated during spermatogenesis if they are inserted into actively transcribed genes. “High and Low” SVA_B–F copies were not enriched in a gene body (Fig. 5B). However, genes with “High and Low” SVA_B–F copies in the antisense direction showed higher expression in spermatogonial stem cells than randomly extracted genes and genes with “Common Low” SVA_B–F copies but lower expression than genes with “Common High” SVA_B–F copies (Fig. 5C). Although approximately half of the “High and Low” SVA_B–F copies were located in the non-genic regions, RNA-seq reads from previously reported undifferentiated spermatogonia (Tan et al. 2020) were more frequently mapped around the non-genic “High and Low” SVA_B–F copies than “Common Low” B–F copies (Fig. 5D). Therefore, genomic regions with the non-genic “High and Low” SVA_B–F copies were potentially transcribed during spermatogenesis. These data suggest that SVAs are subjected to transcription-directed *de novo* DNA methylation during spermatogenesis, and their effectiveness varies among individuals.

**Fig. 5.**
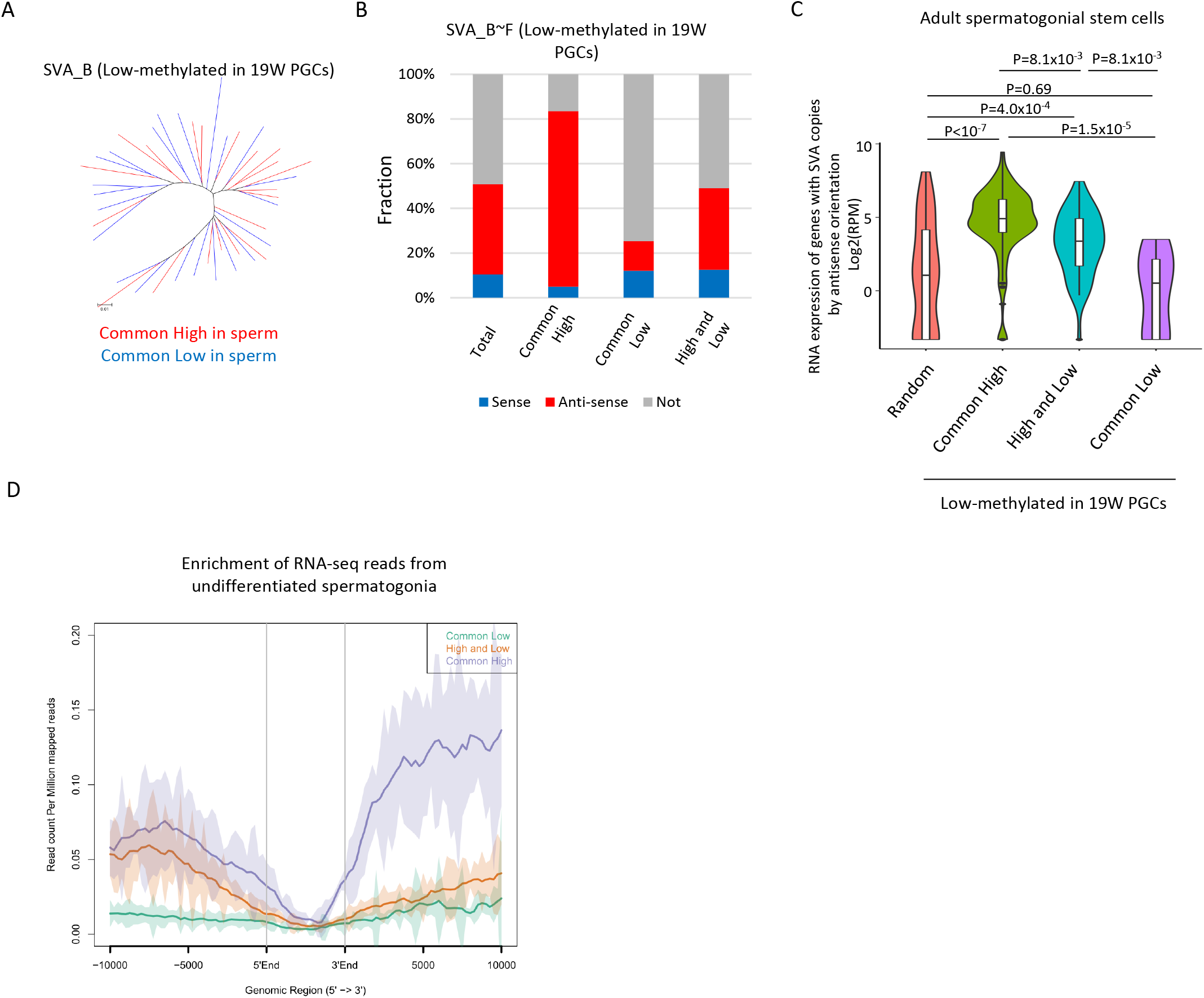
Transciprition associated regulation of DNA methylation of SVA during spermatogenesis (A) Phylogenetic analysis of SVA_B copies low-methylated in male hPGCs at 19 weeks of gestation. SVA_B copies highly methylated by both sperm donors were colored by red, while those hypomethylated by both sperm donors were colored by blue. (B) Bar graphs showing the fraction of SVA_B∼F copies inserted in a gene body. SVA copies were classified by DNA methylation patterns in two sperm donors. Only low methylated SVA copies in male hPGCs at 19 weeks of gestation were used for this analysis. (C) Violin plots showing the expression of genes in adult spermatogonial stem cells 2 (Sohni et al. 2019). Genes were classified according to the DNA methylation status of SVAs inserted in them in the antisense direction. P-value was calculated by Tukey’s test. (D) Enrichment of RNA-seq reads from undifferentiated spermatogonia (Tan et al. 2020) around non-genic SVAs. Only low-methylated SVA copies male hPGCs at 19 weeks of gestation were used for the analysis, and SVA copies were classified by DNA methylation patterns in two sperm donors (common low, high & low and common high).

### SVAs constitute a major source of inter-individual epigenetic variations in sperm

To investigate whether the inter-individual variation of sperm DNA methylation in SVAs was a common phenomenon, we reanalyzed another set of previously reported WGBS data from three sperm donors (Okae et al. 2014). We again observed a significant difference in methylation levels of “High and Low” SVA_B–F copies (Fig. 6A). To further validate this, we performed amplicon-seq of bisulfite PCR products from SVAs (Fig. 6B). For amplicon-seq, we designed three sets of PCR primers in SVAs, and the three PCR products were combined and sequenced using a next-generation sequencer (Fig. 6B). We constructed an amplicon-seq library from five sperm donors and obtained ∼1.7–2.2M read pairs from each sample. More than 90% of full-length SVA_B–F could be analyzed by our SVA amplicon-seq (minimum depth of CpG ≥ 5, analyzed CpG number ≥ 10) (Fig. 6C). Consistent with publicly available sperm WGBS data, “High and Low” SVA_B–F copies showed variation in DNA methylation among individuals (Fig. 6D). Finally, we investigated the impact of SVAs on inter-individual epigenetic variations in sperm. To this end, we identified differentially methylated regions (DMRs) between human sperm donors from previously reported sperm WGBS data (Hammoud et al. 2014). Although DNA methylation profiles between these two donors were highly correlated (Fig. 6E), 2,008 regions were identified as DMRs (donor 1 < donor 2: 332, donor 2 < donor1: 1,676). Out of 1,676 Donor1-specific methylated DMRs, 772 (46.1%) were overlapped with SVAs (Fig. 6F). We also observed differential methylation among individuals in SVA-associated DMRs from our amplicon-seq data (Fig. 6G). Thus, variation in SVA methylation is a major source of the sperm epigenome among human individuals.

**Fig. 6.**
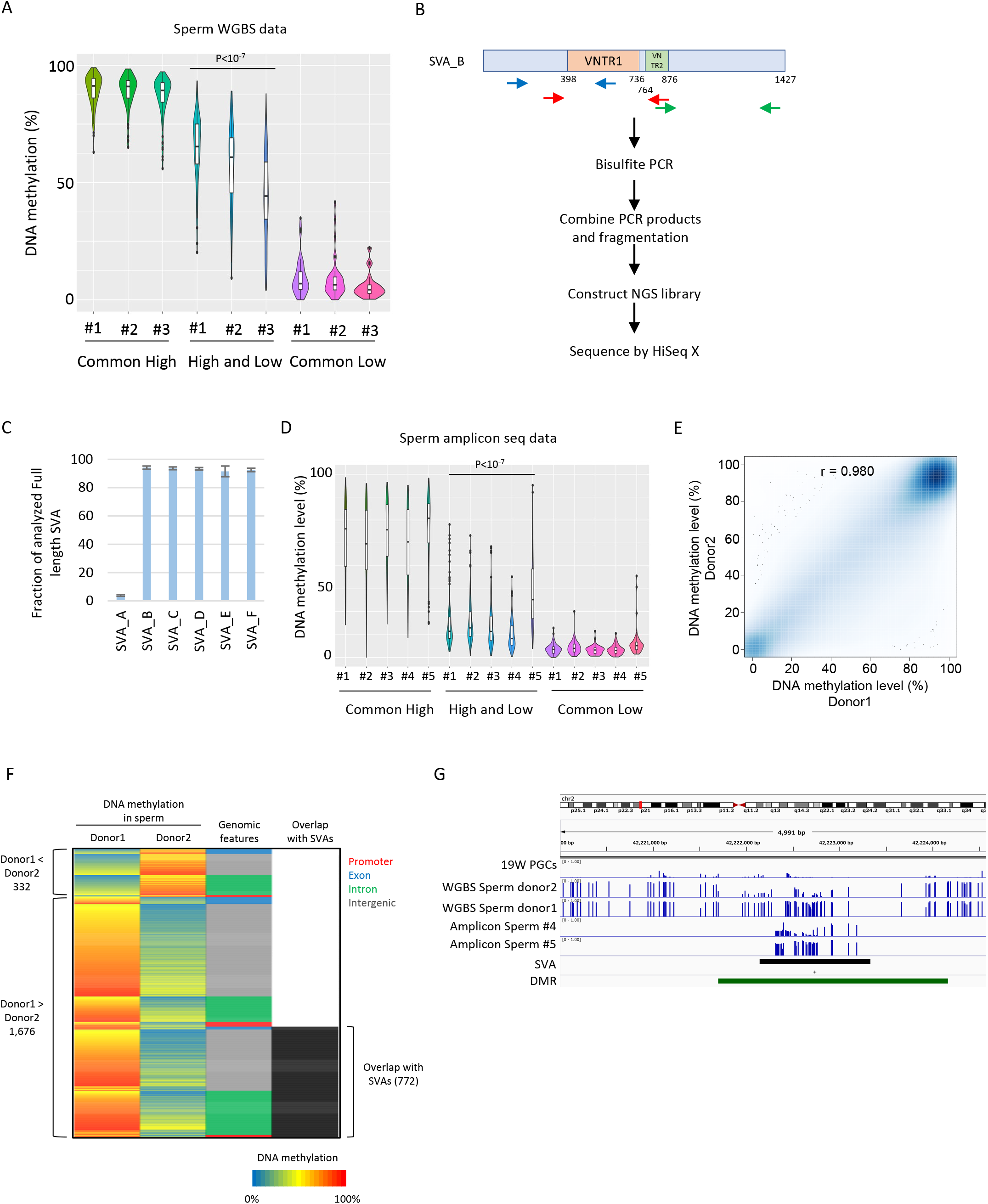
SVAs constitute a major source of inter-individual epigenetic variations in sperm (A) Violin plots showing DNA methylation of SVA copies in previously reported three sperm donors (Okae et al. 2014). Only low-methylated SVA copies in male hPGCs at 19 weeks of gestation were used for the analysis. SVA copies were classified by DNA methylation levels of two sperm donors from *Hammoud et al*. (Hammoud et al. 2014). Donor#1 showed significantly higher DNA methylation levels in “High and Low” SVA copies than other sperm donors. P-value was calculated by Tukey’s test. (B) Scheme of amplicon-seq for analyzing SVA methylation. (C) Bar plots showing the fraction of analyzed full-length SVA copies by amplicon-seq. (D) Violin plots showing DNA methylation levels of SVA copies in five sperm donors from amplicon-seq. Only low-methylated SVA copies in male hPGCs at 19 weeks of gestation were used for the analysis. SVA copies were classified by DNA methylation levels of two sperm donors from *Hammoud et al*.. Donor#5 showed significantly higher DNA methylation levels in “High and Low” SVA copies than other sperm donors. P-value was calculated by Tukey’s test. (E) Scatter plot showing the DNA methylation between sperm donor1 and sperm donor2 from *Hammoud et al*… DNA methylation levels between these two donors were highly correlated. (F) Heatmap showing DNA methylation levels, genomic distribution and overlap with SVAs of DMRs. (G) Representative view of DMRs overlapping with SVA. Black and green boxes represent SVA and DMR, respectively.

## Discussion

In this study, we described the possible molecular mechanism by which retroelements escape from genome-wide DNA demethylation in hPGCs, and how *de novo* DNA methylation is acquired during human spermatogenesis. Our analysis also showed that DNA demethylation resistance in hPGCs frequently occurred in moderately young retroelements such as L1PA, SVA_A, and LTR12. In addition, KRAB-ZFP binding potentially contributed to DNA demethylation resistance of L1s and SVAs. In particular, ZNF257/28 and ZNF649 were associated with DNA demethylation resistance of SVAs and L1s, respectively (Fig. 7). Although it has been reported that ZNF91 binds to VNTR in SVAs and silences SVA expression in embryonic stem cells (ESCs) (Jacobs et al. 2014; Haring et al. 2021), DNA demethylation resistance of SVAs did not correlate with ZNF91 binding, suggesting that a different KRAB-ZFP set is used to suppress SVAs between PGCs and ESCs in humans. Both ZNF257 and ZNF28 bound to VNTR1 (Fig. 2F), and high copy numbers of VNTR1 were correlated with high ZNF257 and ZNF28 enrichment and DNA methylation (Fig. 2H, I). The decreased copy number of VNTR1 after SVA_B emergence (Fig. 2G) may have been necessary for SVAs to escape the silencing mechanisms in hPGCs. We also found that ZNF649 binding was correlated with the DNA demethylation resistance of L1s in hPGCs. Even though our data showed a strong correlation between KRAB-ZFPs and DNA demethylation resistance, direct evidence remains elusive due to the limited availability of human fetal gonads. The PGC-like cells (PGCLCs) *in vitro* derivation systems are thought to be a promising model for investigating the biology of PGCs. Although the successful establishment of human PGCLCs has been reported previously (Sasaki et al. 2015), sufficient DNA demethylation was not observed in human PGCLCs (von Meyenn et al. 2016). Thus, the current human PGCLCs are not a suitable model for investigating the mechanisms of DNA demethylation resistance; further optimizating derivation conditions for human PGCLCs will aid our understanding of retroelement silencing in PGCs.

**Fig. 7.**
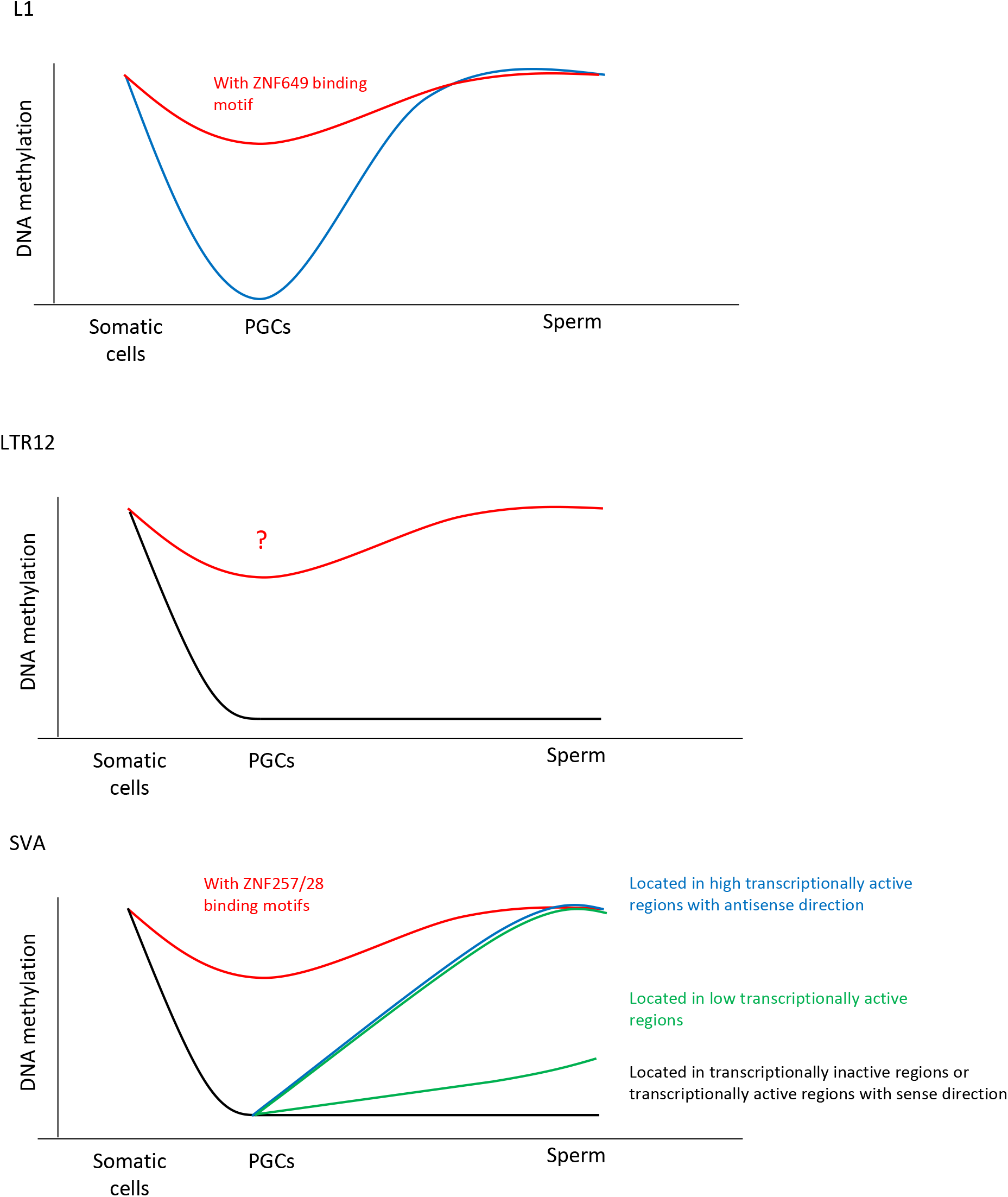
Summary of this study Our data demonstrated the association between KRAB-ZFP binding motifs and DNA demethylation resistance of L1 and SVA. ZNF649 and ZNF257/28 were associated with DNA demethylation resistance of L1 and SVA, respectively. Although we could not find KRAB-ZFPs associated with DNA demethylation resistance of LTR12, phylogenic analysis of LTR12C copies showed that highly methylated copies were genetically separated from hypomethylated copies. Thus, the DNA demethylation resistance of LTR12 is genetically determined, not stochastically or environmentally. The dynamics of DNA methylation during spermatogenesis was largely different among retroelement types. The majority of L1 copies acquired DNA methylation during spermatogenesis, while DNA methylation status of LTR12 in hPGCs tended to be maintained during spermatogenesis. The mode of DNA methylation change in SVAs during spermatogenesis was largely different among copies and individuals. SVA copies located in high transcriptionally active regions acquired DNA methylation during spermatogenesis, while those located in transcriptionally inactive regions maintained a hypomethylation state during spermatogenesis. On the other hand, the degree of DNA methylation in sperm in SVA copies located in low transcriptionally active regions was highly variable among individuals. These results suggest that SVA is methylated by transcription-directed DNA methylation mechanisms during spermatogenesis, and the activity of the mechanisms is highly variable among individuals.

In this study, we also showed that the mode of DNA methylation acquisition during spermatogenesis was vastly different among retroelement types. The majority of L1 copies acquired DNA methylation during spermatogenesis, while LTR12 tended to maintain DNA methylation status in PGCs during spermatogenesis (Fig. 7). L1HS, in which both ZNF93 and ZNF649 could not bind, also acquired DNA methylation during spermatogenesis (Fig. 4B). Other factors may be involved in the DNA methylation of L1 during spermatogenesis. The PIWI-piRNA pathway is responsible for DNA methylation of L1 transposons in mouse male germ cells (Aravin et al. 2007; Carmell et al. 2007; Shoji et al. 2009; Manakov et al. 2015; Kojima-Kita et al. 2016). The PIWI-piRNA pathway might also be functional in humans because mutations in genes involved in the PIWI-piRNA pathway are linked to human male infertility (Gu et al. 2010; Arafat et al. 2017), and the majority of putative piRNAs mapped to transposons at gestational week 20 are derived from L1 (Reznik et al. 2019). Thus, the PIWI-piRNA pathway is a candidate for L1 silencing in human male germ cells. Interestingly, our data demonstrated that the acquisition of DNA methylation of SVAs during spermatogenesis is regulated by their inserted regions, not by their sequences. SVAs inserted in transcriptionally active regions in the antisense direction were targeted by *de novo* DNA methylation during spermatogenesis. It was reported that MIWI2 binds piRNAs and is recruited to nascent transcribed regions that are complementary to piRNAs in mice (Watanabe et al. 2018), then MIWI2 interacting protein SPOCD1 recruits chromatin remodeling complex DNMT3A and DNMT3L to MIWI2 bound genomic regions to induce DNA methylation (Zoch et al. 2020). Thus, nascent transcripts with antisense SVAs could be targeted by the MIWI2/SVA-derived piRNA complex. There are other transcription-directed repetitive element silencing mechanisms, such as the HUSH complex that represses L1s and SVAs (Fukuda et al. 2018; Liu et al. 2018; Robbez-Masson et al. 2018). The HUSH complex targets young, full-length L1s placed within introns of actively transcribed genes (Liu et al. 2018; Fukuda and Shinkai 2020). In addition to the HUSH complex, efficient pericentromeric heterochromatin formation requires transcription of pericentromeric satellite repeats, which stabilizes the SUV39H pericentromeric localization (Johnson et al. 2017; Shirai et al. 2017; Velazquez Camacho et al. 2017). As SUV39H is also involved in retroelement silencing (Bulut-Karslioglu et al. 2014), both the HUSH complex and SUV39H are also candidate factors involved in the transcription-directed DNA methylation of SVAs in human male germ cells. Our data indicated that, in sperm, the degree of DNA methylation of SVAs located in genomic regions with low transcriptionally activity varied among individuals. This result suggests that the effectiveness of transcription-directed *de novo* DNA methylation in human male germ cells varies among individuals. Previous studies have reported that hypermethylation of the promoter regions of *PIWIL2* and *TDRD1*, which are involved in the PIWI-piRNA pathway, is associated with abnormal DNA methylation and male infertility in humans (Heyn et al. 2012). Thus, the functionality of the PIWI-piRNA pathway could be different among individuals, which might also contribute to the epigenetic variation of SVAs in male germ cells. SVAs function as enhancers (Gianfrancesco et al. 2017), and SVA insertions alter the chromatin state around the insertion site (Fukuda et al. 2017). SVA insertions are associated with Fukuyama-type congenital muscular dystrophy and Lynch syndrome (Ostertag et al. 2003; Payer and Burns 2019). Thus, the difference in SVA regulation among individuals may induce changes in gene regulation in male germ cells, the risk of genome instability, and the incidence of diseases among individuals.

## METHODS

### Semen collection

Ejaculates were provided by the patients who visited the Reproduction Center in the Ichikawa General Hospital, Tokyo Dental College. All study participants were briefed about the aims of the study and the parameters to be measured, and consent was obtained. The study was approved by the ethics committees of RIKEN, Tokyo University and Ichikawa General Hospital. Sperm concentration and motility were measured with a computer-assisted image analyzer (C-Men, Compix, Cranberry Township, PA, USA). Human semen was diluted twice with saline, and was layered on 5.0 mL of 20 mM HEPES buffered 90% isotonic Percoll (Cytiba, Uppsala, Sweden) and centrifuged at 400× g for 22 min. The sperm in the sediment was recovered to yield 0.2 mL, and then introduced to the bottom of 2.0 mL of Hanks’ solution to facilitate swim-up. The motile sperm in the upper layer was then recovered.

### Preparation of SVA amplicon-seq

Genomic DNA was subjected to bisulphite-meidated C to U conversion using MethylCode Bisulfite Convrsion Kit (ThermoFisher Scientific), and then used as a template for PCR for 35 cycles with EpiTaq (TAKARA) using the following primer: SVA_1_Fw TTATTGTAATTTTTTTGTTTGATTTTTTTGTTTTAG. SVA_1_Rv AAAAAAACTCCTCACATCCCAAAC. SVA_2_Fw TTAATGTTGTTTAGGTTGGAGTGTAGTG. SVA_2_Rv CAAAAAAACTCCTCACTTCCCAATA. SVA_3_Fw TTTGGGAGGTGTATTTAATAGTTTATTGAGAA. SVA_3_Rv TAAACAAAAATCTCTAATTTTCCTAAACAAAAAACC). The PCR products from 3 set of perimers were combined, were purified by MinElute PCR Purification Kit (QIAGEN), and were fragmented by Picoruptor (Diagenode) for 10 cycles of 30 seconds on, 30 seconds off. Then, amplicon-seq library was constracted by KAPA LTP Library Preparation Kits (KAPA BIOSYSTEMS) and SeqCap Adapter Kit A (Roche). The amplicon-seq libraries were sequenced on a HiSeq X platform (Illumina).

### WGBS and amplicon-seq analysis

#### Quality control, read mapping and calculation of DNA methylation

Low quality bases and adaptor sequences were trimmed by Trim Galore version 0.3.7 (http://www.bioinformatics.babraham.ac.uk/projects/trim_galore/). Then trimmed reads were aligned to the hg19 genome by Bismark v0.14.1 (Krueger and Andrews 2011). The methylation level of each CpG site was calculated as follows: (number of methylated reads/number of total reads). Only CpG sites with at least five reads were used for all analysis. Only nearly-full length retroelements, which contain at least 90% portion of consensuse sequence, were used for DNA methylation analysis of retroelements. The information of retroelements was obtained from UCSC genome browser (http://genome.ucsc.edu/). For DNA methylation analysis of retroelements, we used retroelements containing at least 10 CpG sites of which read depth is at least five reads. The methylation level of each retroelement copy was calculated by averaging the methylation levels of the CpG sites within the copy.

#### Classification of retroelement copy according to DNA methylation level

Retroelement copies were classfied by according to their DNA methylation levels as follows: Low < 20%, 20% <= Medium < 60%, High >= 60%.

#### Association of KRAB-ZFP peaks, binding motifs and retroelement

We obtained peak regions of each KRAB-ZFP, which were previously reported (Imbeault et al. 2017), from gene expression omunibus GSE78099. Overlap of KRAB-ZFP peak and retroelement copy was investigated by bedtools v2.15.0 (Quinlan and Hall 2010). The binding motif of each KRAB-ZFP was predicted by the findMotifsGenome.pl program in Homer v4.8.3 (Heinz et al. 2010). The KRAB-ZFPs binding motifs along retroelement copies were searched by FIMO (Grant et al. 2011). We used predicted motif sites with a q-value of 0.00005 or less for ZNF257 and ZNF28 and with a q-value of 0.05 or less for ZNF93 and ZNF649 in this study.

#### DMR identification

DMR candidates were identified using the ‘Commet’ command in BisulFighter (Saito et al. 2014). To enhance the confidence of DMR call, we calculated the average methylation levels of the candidates using CpG sites with >=5 reads in both sperm donors, and among the candidates, those containing >=10 successive analyzable CpG sites and showing a >=40% methylation difference were determined as DMRs.

### Phylogenetic analysis of retroelement copies

The evolutionary history was inferred by using the Maximu Liklihood method based on the Tamura-Nei model (Tamura and Nei 1993). Initial tree(s) for the heuristic search were obtained by applying the Neighbor-Joining method to a matrix of pairwise distances estimated using the Maximum Composite Likelihood (MCL) approach. The tree is drawn to scale, with branch lengths measured in the number of substitutions per site. There were a total of 10153 positions in the final dataset. Evolutionary analyses were conducted in MEGA6 (Tamura et al. 2013).

### RNA-seq analysis

We reanalyzed previously reported single-cell RNA-seq data in testis (Sohni et al. 2019). Read counts data of genes and cell type annotation of each cell were obtained from Gene Expression Omnibus under accession number GSE124263. Reads per million mapped reads (RPM) for genes were calculated in each cell. We used the average RPM of spermatogonial stem cells 2 in Fig. 5C. We also reanalyzed previously reporeted RNA-seq data from undifferentiated spermatogonia (Tan et al. 2020), which is deposited in Gene Expression Omunibus under accession number GSE144085. Low quality bases and adaptor sequences were trimmed by Trim Galore version 0.3.7. Then, trimmed reads were aligned to the hg19 genome by Bowtie v0.12.7 with ?m 1. Enrichment of RNA-seq reads around SVAs were visualized by ngsplot. RNA expression levels of ZNF93 and ZNF649 in hPGCs (Fig. 3F) were obtained from GSE63392 (Gkountela et al. 2015).

### Visualization of NGS data

The Integrative Genomics Viewer (IGV) (Robinson et al. 2011)was used to visualize NGS data. Enrichment of RNA-seq reads and KRAB-ZFPs was visualized by ngsplot (Shen et al. 2014). Scatter plot and violin plot analysis were performed by ggplot2 package in R.

## DATA ACCESS

All reads from amplicon-seq in this study have been submitted to Gene expression omnibus under accession number GSE174562.

## ACKNOWLEDGEMENTS

We thank the staff of the Support Unit for Bio-Material Analysis (BMA) at RIKEN Center for Brain Science (CBS) Research Resources Division (RRD) for NGS library construction. This research was supported by the Special Postdoctoral Researcher (SPDR) Program of RIKEN to KF, the Japan Ministry of Education, Culture, Sports, Science and Technology Grant-in-Aid for Scientific Research (18H05530, 18H03991) to YS and RIKEN internal research funds to YS.

## AUTHOR CONTRIBUTIONS

Conceptualization: YS, KF, KI. Sample collection: YM, SK, YO.

